# FERROPTOSIS GENE SIGNATURES REVEAL DISTINCT REGULATORY LANDSCAPES IN GASTRIC ADENOCARCINOMA AND OTHER TISSUES

**DOI:** 10.1101/2025.09.11.675514

**Authors:** Sérgio Augusto Antunes Ramos, Thaynara Pereira de Abreu, Marcos da Conceição, Valéria Cristiane Santos da Silva, Diego Pereira, Daniel de Souza Avelar, Jéssica Manoelli Costa da Silva, Ronald Matheus da Silva Mourão, Rubem Ferreira da Silva, Williams Fernandes Barra, Samir Mansour Casseb, Andrea Kely Campos Ribeiro dos Santos, Samia Demachki, Geraldo Ishak, Fabiano Cordeiro Moreira, Rommel Mario Rodriguez Burbano, Paulo Pimentel de Assumpção

## Abstract

**Background:** Gastric adenocarcinoma (GAC) remains one of the most lethal malignancies worldwide, with late-stage diagnosis and limited therapeutic options. Ferroptosis, a regulated form of cell death driven by iron-dependent lipid peroxidation, has emerged as a promising target for overcoming tumor resistance mechanisms. This study aimed to characterize the transcriptional landscape of ferroptosis-related genes in GAC, comparing tumor, peritumoral, metaplastic, and normal gastric tissues.

**Methods:** RNA-Seq was performed on 385 biopsied samples from patients treated at the João de Barros Barreto University Hospital. Differential expression analysis was conducted using DESeq2, and genes related to ferroptosis were identified based on FerrDb V2 annotations. Visualization included volcano plots, DAPC clustering, heatmaps, and gene dominance scoring.

**Results:** GAC samples showed a distinct ferroptotic expression signature, with simultaneous upregulation of key promoters (e.g., *CDKN2A, NOX4, EGFR, IL6*) and suppressors (e.g., *HSPB1, SCD, NUPR1, GDF15*). Notably, the tumor tissue exhibited a net dominance of ferroptosis-inhibitory genes, suggesting an adaptive response to oxidative stress. Adjacent tissues showed partial overlap with tumor profiles, while metaplastic tissue displayed a hybrid signature with selective suppression of ferroptosis. Normal mucosa exhibited dominant expression of promoters, contrasting with the tumor’s anti-ferroptotic phenotype.

**Conclusion:** The transcriptional heterogeneity and regulatory imbalance of ferroptosis-related genes in GAC support its role as a potential therapeutic axis. These findings provide molecular insights for biomarker discovery and ferroptosis-targeted strategies in gastric cancer.

## 1 Introduction

Gastric cancer is the fifth most common type of cancer and the fourth with the highest mortality rate worldwide, constituting one of the most prevalent malignancies. According to data from GLOBOCAN, approximately 968,000 new cases and 660,000 deaths attributed to the disease were recorded^1^. In Brazil, the estimated number for each year of the 2023-2025 triennium is 21,480 new cases. Although statistics indicate a downward trend in global incidence, diagnosis still occurs largely at advanced stages, which limits the effectiveness of conventional therapeutic approaches.

In this context, the search for more effective therapies has gained notoriety, especially among those capable of evading tumor resistance mechanisms, including the inactivation of cell death pathways, particularly apoptosis and necrosis. Assessing how alternative death pathways, such as ferroptosis, can contribute to containing tumor growth has gained focus in recent years^2^.

Ferroptosis^3^ is a form of cell death caused by iron-dependent lipid peroxidation and reactive oxygen species (ROS). Unlike apoptosis and necrosis, it is characterized by redox imbalance, elevated ROS levels, and disruption of the mitochondrial outer membrane^3^. Ferroptosis can occur through two main pathways: the external or transporter-dependent pathway and the internal or enzyme-regulated pathway^4^. It can be induced through drug intervention for the treatment of specific diseases such as Alzheimer’s, Parkinson’s, and cancer.

In the carcinogenic context, cell death through ferroptosis can be used in a targeted manner as an inhibitor of tumor progression, since extracellular iron supplementation makes them more vulnerable to ferroptosis inducers through the inactivation of glutathione peroxidase 4 (*GPX4*), an enzyme that functions as a central regulator of ferroptosis, generating accumulation of phospholipid hydroperoxides (POOH), causing damage to the cell membrane and leading to ferroptotic death^5^.

Cancer therapeutics using ferroptosis may become a promising treatment avenue, since high iron levels in tumor cells can be induced to produce large amounts of ROS and POOH, eliminating tumor cells^5^. This work aims to investigate the transcriptional landscape of genes linked to ferroptosis in gastric adenocarcinomas, differentiating expression profiles between tumor and non-tumor tissues, based on the promoter or preventative function of these genes in inducing cell death.

## 2 Material and methods

### 2.1) Ethical aspects and sampling

The patients participating in the study were treated at the João de Barros Barreto University Hospital (HUJBB). Sample collection was performed only after the patients signed an Informed Consent Form (ICF) and were informed of the risks and benefits of the study. The present study was approved by the hospital’s Research Ethics Committee (CAAE No. 47580121.9.0000.5634) for sample collection, which was stored in RNAlater and preserved at −80°C. Participant recruitment and sample collection were carried out between July 2, 2022, and July 6, 2023.

A total of 385 tumor tissue samples were collected through gastric biopsies from patients diagnosed with gastric adenocarcinoma (GAC). The samples were divided into four groups: 1. GAC, 156; 2. Metaplasia, 20; 3. Peritumoral tissue (PTT), 186; and 4. Normal gastric mucosa, 23.

### 2.2) RNA extraction and quality assessment

Tumor tissues from each sample were macerated for total RNA extraction using TRIzol reagent (Thermo Fisher Scientific), according to the manufacturer’s instructions. Quality assessment was performed by quantifying RNA using a Qubit 4.0 Fluorometer (Thermo Fisher Scientific) and assessing integrity using agarose gel electrophoresis (1g) with Safer Dye (4µ) (QIAGEN, Germany).

### 2.3) Library preparation and NGS sequencing

Libraries for sequencing were constructed using the TruSeq Stranded Total RNA Library Prep Kit with Ribo-Zero Gold (Illumina, USA), according to the manufacturer’s instructions, using 1 μg of total RNA per sample in a total volume of 11 μL. Subsequently, the samples were evaluated for the integrity of the generated cDNA fragments using the 2200 TapeStation system (Agilent Technologies AG, Basel, Switzerland). Next generations sequencing was performed using the paired-end approach on the NextSeq® 500 platform (Illumina®, USA) with the NextSeq® 500 MID Output 2 x 75 bp, 150 cycles kit (Illumina, USA), according to the manufacturer’s instructions.

### 2.4) Read quality analysis

After sequencing, the quality of RNA-Seq reads was assessed using the FASTQC program^6^. They then underwent a trimming step, removing low-quality bases and adapters using the FASTP software^7^. A quality value (QV) threshold greater than 15 was applied, and the files were then forwarded to the NEXTFLOW pipeline^8^.

### 2.5) Quantification and alignment of human transcripts

In the NEXTFLOW pipeline^8^, the trimmed data were aligned against the human genome using the SALMON^9^ program, and their transcripts were quantified using the STAR^10^ software, using hg v38 (www.ensembl.org) as the reference index and GENCODE v.42 (www.gencodegenes.org) as the annotation object.

### 2.6) Differential Analysis and Data Normalization

After the previous step, the data were imported into the RStudio^11^ platform (version 4.4) using the TXImport package^12^. Differential analysis was performed using the DESeq2^13^ library, whose parameters for selecting differentially expressed genes (DEGs) were |log2(fold-change)| > 1 and adjusted p-value < 0.05. The design used in the DESeq function of this step involved tumor samples (GAC) and three comparator tissues: adjacent tissue, metaplasia, and normal gastric mucosa. The data were normalized using the model-based variance-stabilizing transformation (VST)^14^ method.

To visualize the clustering of the four sample types, Discriminant Analysis of Principal Components (DAPC) was performed, which is illustrated in a 3D scatter plot generated by the Plotly^15^ package.

### 2.7) Ferroptosis gene selection

Genes related to the ferroptosis pathway were filtered using gene lists documented in the FerrDB V2^16^ database, a repository dedicated to providing diverse information on ferroptosis regulators. Two lists were downloaded: one containing genes that promote ferroptosis (called Promoters) and one containing genes that prevent the pathway (called Preventers). Only signatures tested in humans or humans/mice remained in the analysis.

The search for these genes in the RNA-Seq data was performed using the Dplyr^17^ data manipulation package in the RStudio^11^ environment. The graphical data visualization language was provided by the Ggplot2^18^ package, which generated volcano and bar plots.

The heatmaps were processed using the ComplexHeatmap package^19^, in which the data were z-score transformed and several annotations were added to reflect all the clinical characteristics of the presented data. To better visualize the distribution of samples by tissue type and the regulatory nature of the genes, the “row_split” parameter was used in the dendrograms to group samples of the same type.

The “gene type dominance” annotation was added to generate a scatter plot at the top of the heatmaps, indicating the difference in mean expression between promoter and preventer genes for each sample. Positive values indicate a greater tendency for promoter gene expression to contribute to ferroptosis; negative values indicate a greater tendency for preventer genes, providing a global overview of what each tissue type expresses in relation to ferroptosis regulation.

## 3 Results

Differential analysis revealed three distinct scenarios, in which a consistent pattern of gene dysregulation is observed across the analyzed groups. The volcano plots illustrated in Figure 1 highlight the set of ferroptosis-related genes previously filtered from FerrDb v2. In the comparison between GAC and PTT (Figure 1A), several ferroptosis-related genes were differentially upregulated in the tumor. Among these are the ferroptosis promoters *CDKN2A, NOX4, YAP1*, and *EGFR*, while others that act to repress ferroptosis were *SCD, TYRO3*, and *SREBF1*.

**Figure 1.**
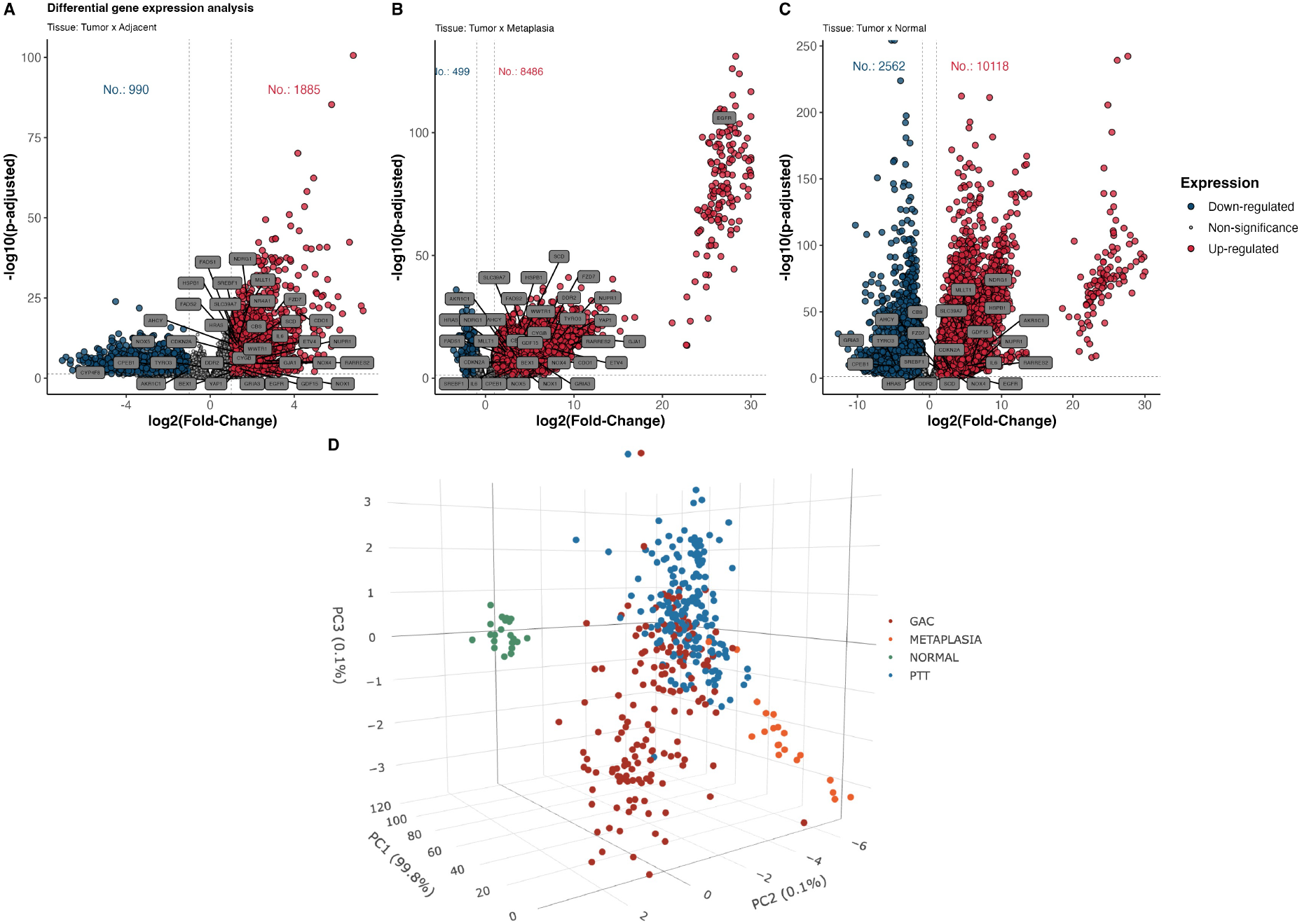
Differential expression analysis between GAC and PTT (A), metaplasia (B), and normal gastric mucosa (C) presented by volcano plots in which rectangular labels indicate genes linked to ferroptosis and red dots symbolize upregulation, blue downregulation, and gray dots, with no statistical significance. In D, 3D scatter plot of DAPC for all study groups.

Between GAC and metaplasia (Figure 1B), an even more unbalanced profile was observed, as all ferroptotic genes were upregulated in the tumor, including *EGFR, NOX-1, −4, −5*, and *HSPB1*. This characteristic indicates an important finding for raising hypotheses about the regulatory landscape of ferroptosis in metaplastic cells.

In the comparison between GAC and normal mucosa (Figure 1C), the contrast was even more pronounced, as 10,118 genes were upregulated in the tumor, compared to only 2,562 downregulated. This discrepancy stems from the biological differences between the two tissues and the imbalance in the number of samples from each. In this scenario, both pro- and anti-ferroptosis genes were found in the tumor and normal tissue.

DAPC highlights the sample clusters, with GAC and PTT overlapping and partially clustered with the metaplastic sample, with the group of normal mucosa samples in the background. The overlap observed between the tumor and adjacent tissue reveals a similar expression profile shared by both tissues, while the metaplastic tissue presents some distance from these groups. However, the controls show that there are indeed significant differences between the normal and altered expression profiles.

In the expression profile overview, Figure 2 summarizes the transcriptional expression of the 35 ferroptosis-related genes identified in the data, distributed across the tissues. The genes were categorized according to their functional role as ferroptosis promoters or preventers, as curated by FerrDb v2. A highly heterogeneous expression pattern is noted across genes and samples. Ferroptosis-promoting signatures, such as *CDKN2A, CYGB, EGFR, YAP1, WWTR1, DDR2, GJA1*, and *IL6*, as well as those that prevent this pathway, such as *GDF15, AHCY, SCD, HSPB1, RARRES2, CBS, NUPR1*, and *NR4A1*, exhibited high expression (positive z-score) predominantly in tumor samples when compared to other tissues.

**Figure 2.**
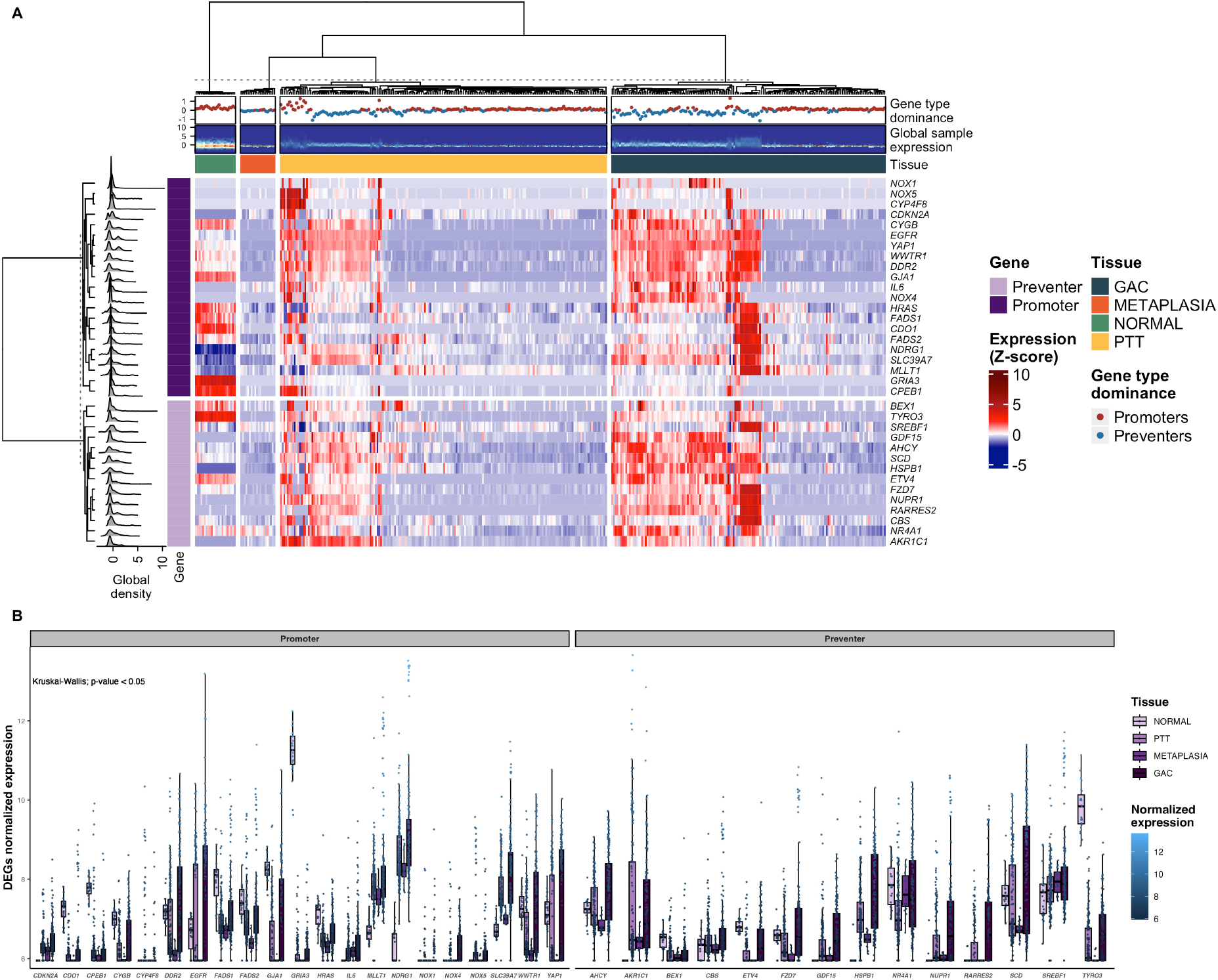
Heatmap showing the z-score expression of ferroptosis genes across all sequenced tissue types, where blue tones reflect low expression, while red tones, high expression. The left side contains two annotations: one with a density plot (gray) whose peaks indicate the value at which most samples showed expression, and the second (light/dark purple) represents the regulatory nature of each gene (right). At the top, from top to bottom, is a scatter plot (red/blue dots) indicating the average dominance of the gene type in each sample; in the middle, a mini-heatmap highlights regions of greater or lesser distribution of values; finally, each bar designates a sample type (A). Boxplot with the normalized expressions for the 35 ferroptosis genes found in each tissue (B).

However, the gene type dominance plot (Figure 2A top) shows a strong tendency toward higher expression of Preventer genes (blue dots) in GAC, suggesting that the tumor is leading to selective activation of antiferroptotic pathways, which may be accompanied by repression of inhibitory mechanisms, a possible reflection of tumor adaptation to oxidative stress, metabolic remodeling, and death evasion (cellular immortalization). In contrast, a greater tendency for the average expression of Promoter genes (red dots) is observed across the PTT samples, which may be linked to the canonical activation of this death pathway as part of the natural physiological process of this tissue. Furthermore, the dominance of Promoters in normal tissue reinforces this finding (red dots in all samples).

In the context of metaplastic tissue, curiously, a few gene signatures showed slightly elevated expression in some samples, such as the Preventers *SREBF1, SCD*, and *NR4A1*, and the Promoters *NDRG1* and *MLLT1* (Figure 2B). The dominance scatter plot shows a tendency toward genes that prevent ferroptosis, perhaps because this tissue is constantly undergoing genetic reprogramming to adapt to new conditions and, therefore, prevents the activation of death pathways.

The left-hand annotation of the heatmap (global density) reveals that, although some genes exhibit broad expression across all tissues (*IL6, CDKN2A, EGFR*), most samples exhibit downregulation restricted to subsets, suggesting a possible association with specific molecular subtypes or microenvironments (Figure 2A). The analysis of these differences is being investigated in a parallel study, which investigates the influence of clinical data on therapeutic management (neoadjuvant therapy), the presence of *Helicobacter pylori*, among others.

When comparing gene expression between GAC and normal tissue, several genes were observed to have reduced expression levels in the latter, in contrast to increased expression in the tumor. This pattern was evident for *AKR1C1, CDKN2A, GDF15, HSPB1, IL6, NOX4, NUPR1*, and *RARRES2* (Figure 3). These genes are related to different cellular processes, including oxidative metabolism (*AKR1C1, NOX4*), cell cycle regulation (*CDKN2A*), stress response (*HSPB1, NUPR1*), inflammatory signaling (*IL6*), growth factors (*GDF15*), and microenvironmental modulation (*RARRES2*). The observed profile reinforces that these genes present clear differential expression between tumor and normal tissue, characterizing them as part of the set of genes regulated in GAC, which raises hypotheses of possible biomarkers of ferroptosis in gastric cancer (Figure 3).

**Figure 3.**
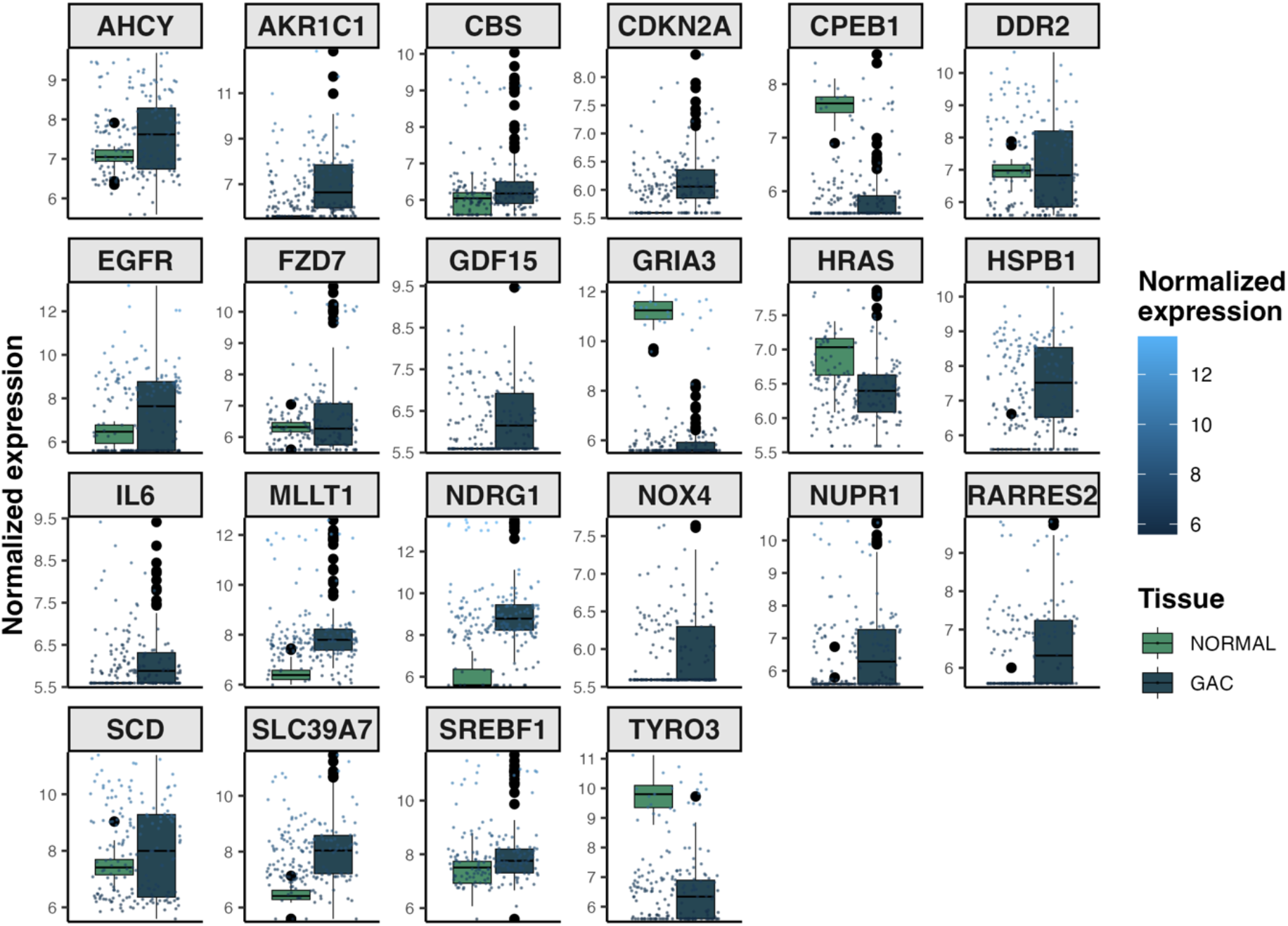
Normalized expression of differentially expressed genes between GAC and normal gastric mucosa. Each panel displays a gene and boxplots accompanied by individual points, representing the biological samples, highlighting intra- and intergroup variation.

Taken together, these findings indicate that the expression profile of ferroptosis genes clearly distinguishes tumor from non-tumor tissues, potentially providing insights for molecular stratification and identification of novel therapeutic targets. Furthermore, this transcriptional landscape points to robust transcriptional reprogramming in the tumor context, with selective activation of pro-ferroptosis pathways and inhibition of preventive genes, possibly reflecting adaptive or compensatory mechanisms of tumor cells to oxidative stress.

## 4 Discussion

### 4.1 Differential activation of the ferroptotic pathway in gastric tumors

Differential expression analysis revealed a robust pattern of gene dysregulation between GAC and non-tumor tissues, with a focus on genes related to ferroptosis. The results indicate that, although both promoters and inhibitors of this pathway are expressed in tumors, there is a functional predominance of anti-ferroptotic mechanisms in GAC. This profile may reflect a metabolic adaptation of the tumor to the high burden of oxidative stress, which corroborates previous findings showing that cancer cells develop mechanisms to suppress ferroptosis as a means of evading cell death^20,21^.

Genes such as *HSPB1, SCD, NUPR1*, and *GDF15*, which act as ferroptosis suppressors, were highly expressed in tumor samples. The *HSPB1* gene, for example, negatively regulates iron entry and lipid peroxidation, and is often associated with resistance to heat stress and tumor survival^22^. *SCD* catalyzes the synthesis of monounsaturated fatty acids, whose incorporation into membrane phospholipids reduces susceptibility to lipid peroxidation, a central step in ferroptosis^23^.

### 4.2 Selective suppression of ferroptosis promoters in adjacent tissues

The partial overlap between the transcriptional profiles of GAC and PTT, evidenced by DAPC analysis, suggests that apparently non-cancerous tissues adjacent to the tumor already exhibit transcriptional alterations consistent with a premalignant state. This finding is consistent with the concept of the cancerization field effect, according to which molecular alterations precede visible histological changes^24–26^.

Interestingly, ferroptosis-promoting genes, such as *EGFR, YAP1, WWTR1*, and *CDKN2A*, showed increased expression in GAC but were also present at moderate levels in adjacent tissues. *EGFR*, in addition to its classical function in proliferation, has been implicated in sensitivity to ferroptosis in several solid tumors, being a potential pharmacological target for combination with ferroptotic inducers^5^.

### 4.3 Metaplastic tissue: a hybrid and adaptive profile

The metaplastic tissue exhibited an intermediate expression pattern, characterized by a punctual elevation of preventive genes such as *NR4A1, SCD*, and *SREBF1*. This profile may indicate a state of adaptive reprogramming, in which ferroptosis is transiently suppressed to preserve cellular integrity in the face of adverse conditions, such as chronic inflammation or *H. pylori* infection^27^. The literature on this topic is scarce; one recent study showed significant differential expression between gastric metaplastic and normal tissue involving 17 ferroptosis-linked genes^28^. However, the study has sample size limitations (Metaplastic: 8, Normal: 16) and contained a gene set that differs significantly from that shown in our study.

The expression of *NDRG1*, a known regulator of cellular differentiation with a controversial role in ferroptosis, was also observed in this tissue. Studies suggest that context-dependent regulation of ferroptosis may be critical in transitional states such as gastric intestinal metaplasia^29^.

### 4.4 Transcriptional contrast with normal gastric mucosa

The comparison between GAC and normal controls revealed the highest number of differentially expressed genes, which is expected given the distinct histological and functional nature of these tissues. Among the most highly expressed genes in tumors are CDKN2A, IL6, *AKR1C1*, and, notably, *NOX4*. While *CDKN2A* and *IL6* are already widely recognized for their involvement in cell cycle regulation and tumor inflammation, respectively, *NOX* family products are emerging as critical modulators of neoplastic progression, particularly through the localized generation of reactive oxygen species (ROS) in the cell membrane^30,31^.

Among these, *NOX4* stands out for its ability to sustain an invasive phenotype through the activation of signal transduction pathways associated with epithelial-mesenchymal transition (EMT), expression of metalloproteinases, and remodeling of the extracellular matrix. Furthermore, its activity is strongly associated with the formation of invadopodia^31^, cellular structures specialized in the degradation of the extracellular environment and essential for tumor invasion and metastasis.

This function is enhanced by the production of ROS, which not only acts as an intracellular second messenger but also amplifies the signaling of soluble factors such as *TGF-β* and *EGF*, promoting greater adhesion, motility, and proteolytic capacity of tumor cells. The overexpression of *NOX4* in GAC, therefore, may reflect a strategic adaptation of the tumor to sustain its aggressiveness and invasive capacity, positioning this gene as a potential prognostic biomarker or therapeutic target in anti-invasive interventions^31^.

The IL6 gene, a key cytokine in tumor inflammation, has been widely implicated in promoting a pro-tumor microenvironment, in addition to modulating cell death pathways, including ferroptosis, via activation of *STAT3* and *NF-κB*^32^.

### 4.5 Translational and therapeutic implications

Mapping the ferroptotic expression profile in GAC offers important clues for molecular stratification and the development of targeted therapies. Overexpression of ferroptosis-inhibitory genes can be explored as a therapeutic target, as sensitizing these cells to ferroptotic death represents a promising approach in gastric cancer resistant to conventional chemotherapy^33^.

Furthermore, the identification of genes such as *RARRES2, NUPR1*, and *GDF15* as differentially expressed may indicate biomarkers for prognosis or therapeutic response, although these findings require functional validation *in vitro* and *in vivo*.

## 5 Conclusion

In this study, we investigated the expression profile of ferroptosis-related genes in gastric adenocarcinoma samples, including normal mucosa, tumor-adjacent tissue, and metaplastic tissue. Differential analysis revealed significant alterations in key genes involved in the regulation of oxidative stress, cell death, and tumor progression, notably the overexpression of suppressors such as *HSPB1* and *SCD*, and promoters such as *NOX4, IL6*, and *CDKN2A*.

The heterogeneity in expression patterns across tissues and the predominance of ferroptosis-regulating genes in the tumor microenvironment reinforce the role of this process as a relevant functional axis in gastric cancer. These findings not only contribute to the understanding of the molecular mechanisms associated with tumor pathogenesis but also support the identification of novel therapeutic targets and biomarkers with potential clinical application.

## Author Contributions

**Conceptualization:** Sérgio Augusto Antunes Ramos, Fabiano Cordeiro Moreira, Paulo Pimentel de Assumpção; **Data curation:** Sérgio Augusto Antunes Ramos, Ronald Matheus da Silva Mourão, Daniel de Souza Avelar, Fabiano Cordeiro Moreira; **Formal analysis:** Sérgio Augusto Antunes Ramos, Ronald Matheus da Silva Mourão, Fabiano Cordeiro Moreira; **Funding acquisition:** Fabiano Cordeiro Moreira, Geraldo Ishak, Samia Demachki, Samir Mansour Casseb, Paulo Pimentel de Assumpção, Rommel Mario Rodriguez Burbano, Williams Fernandes Barra; **Investigation:** Sérgio Augusto Antunes Ramos, Thaynara Pereira de Abreu, Fabiano Cordeiro Moreira, Samir Mansour Casseb, Paulo Pimentel de Assumpção; **Methodology:** Sérgio Augusto Antunes Ramos, Valéria Cristiane Santos da Silva, Diego Pereira, Marcos da Conceição, Daniel de Souza Avelar, Jéssica Manoelli Costa da Silva, Ronald Matheus da Silva Mourão, Rubem Ferreira da Silva, Samia Demachki, Andrea Kely Campos Ribeiro dos Santos, Samir Mansour Casseb, Fabiano Cordeiro Moreira; **Sample Acquisition:** Geraldo Ishak, Paulo Pimentel de Assumpção, Rommel Mario Rodriguez Burbano, Williams Fernandes Barra; **Supervision:** Fabiano Cordeiro Moreira, Samir Mansour Casseb, Paulo Pimentel de Assumpção, Rommel Mario Rodriguez Burbano; **Project Administration:** Fabiano Cordeiro Moreira, Samia Demachki, Samir Mansour Casseb, Paulo Pimentel de Assumpção, Rommel Mario Rodriguez Burbano; **Validation:** Samia Demachki, Williams Fernandes Barra; **Visualization:** Sérgio Augusto Antunes Ramos; **Writing – original draft:** Sérgio Augusto Antunes Ramos, Diego Pereira, Fabiano Cordeiro Moreira; **Writing – review & editing:** Sérgio Augusto Antunes Ramos, Thaynara Pereira de Abreu, Samir Mansour Casseb, Paulo Pimentel de Assumpção, Fabiano Cordeiro Moreira.

## Acknowledgment

The authors would like to thank the Oncology Research Center, the Human and Medical Genetics Laboratory, and the Anatomical Pathology Laboratory at João de Barros Barreto University Hospital (HUJBB – UFPA) for their invaluable technical and laboratory support. Our gratitude also goes to the High-Performance Computing Center (CCAD) at the Federal University of Pará for access to the Apollo 2000 cluster, which was crucial for our analyses.

## Data availability statement

The original contributions presented in the study are included in the article. Further inquiries can be directed to the corresponding author.

